# Differential time-restricted sensitivity of enveloped viruses to Sec61 translocon blockade

**DOI:** 10.1101/2025.02.03.636181

**Authors:** Lucy Eke, Belinda Hall, Lucy Thorne, Katja Ebert, Lauren Kerfoot, Greg Towers, Rachel Simmonds, Gill Elliott

## Abstract

The morphogenesis of enveloped viruses relies on the trafficking of transmembrane proteins through the secretory pathway to sites of virus envelopment. The first step in this pathway, their translocation into the endoplasmic reticulum, is therefore an attractive target for broad-spectrum intervention. Here, we tested if blockade of the Sec61 translocon by the *Mycobacterium ulcerans* exotoxin mycolactone, a potent inhibitor of Sec61, could block the production of virus glycoproteins and subsequent production of infectious virus from a range of human enveloped viruses: the DNA virus herpes simplex virus 1 (HSV1), and the RNA viruses, respiratory syncytial virus (RSV), influenza A virus (IAV), SARS coronavirus 2 (SARS CoV2) and Zika virus (ZIKV). In line with known translocation mechanisms, mycolactone blocked *in vitro* translocation and ectopic expression of type I transmembrane proteins but not type III, multipass or cytosolic proteins. Translocation of the type II protein RSV G was also blocked and although ectopically expressed G protein was detected, it was not glycosylated. Pretreatment of cells with mycolactone also blocked the synthesis of type I transmembrane proteins in infected cells and either the synthesis or glycosylation of type II transmembrane proteins, and the production of progeny from all viruses tested, while having no effect on virus entry or downstream synthesis of cytosolic proteins. While mycolactone treatment of HSV1 infected cells at various times after infection resulted in the immediate inhibition of virus production at the point of addition, IAV, RSV and ZIKV became resistant to the action of mycolactone surprisingly early in infection, and before virus glycoprotein synthesis was even detectable or virus production had begun. We therefore conclude that although inhibition of the translocation of virus transmembrane proteins through the Sec61 translocon can in principle block virus production, the morphogenesis of many enveloped RNA viruses requires only limited amounts of envelope proteins for successful propagation, providing novel insight into the biology of these viruses.

**Author Summary:** Many circulating human pathogens are enveloped viruses that all use the cellular secretory pathway to target their envelope proteins to cellular sites of virus particle assembly. The potential to target this pathway could therefore offer a novel broad-spectrum therapy for existing, emerging and as yet unknown human pathogens. Here we have targeted the initial step in this pathway using a highly potent Sec61 inhibitor, mycolactone, to carry out the first comprehensive assessment of translocation disruption on a range of enveloped human viruses from different virus families, including herpes simplex virus, influenza A virus and SARS-CoV2. Our results have shown that Sec61 inhibition blocks the onward trafficking of many virus envelope proteins that are essential to produce infectious virus at assembly sites. However, unexpectedly, we found that several of the viruses were resistant to the effects of this toxin when it was added early in infection, indicating that the synthesis of these essential virus proteins occurs earlier in infection than previously recognised. Hence, while this approach may not be suitable as a broad intervention strategy, it has revealed new information on virus biology and provides us with a novel tool for exploring a wide range of enveloped viruses.

## Introduction

Many antiviral drugs in use today inhibit virus replication directly by targeting virus components from specific viruses, with points of intervention at virus entry, uncoating, genome replication or virus assembly and/or release [1]. By contrast, host-directed antivirals offer the possibility of broad-spectrum, pan-viral activity because multiple viruses require the same cellular machinery for their replication and assembly [2]. The few approved broad-spectrum host-directed antivirals that are in use include interferons which modulate the host innate immune response [3] and drugs which alter the endosomal environment during virus entry [4]. More specific host targeting often focuses on inhibition of entry receptors and co-receptors, such as maraviroc which targets the co-receptor for HIV infection, CCR5 [5]. A possible broad-spectrum point of intervention for enveloped viruses, which include many circulating human pathogens, is the translocation of envelope transmembrane proteins into the endoplasmic reticulum (ER), an essential process for the morphogenesis of these viruses. Because envelope proteins perform dual actions in virus infection - virus attachment and entry into cells at the start of infection, and recruitment of internal virus structures to membranes during virus morphogenesis – disrupting their biosynthesis would not only block virus propagation but also provide novel information on virus morphogenesis pathways.

Transmembrane proteins fall into several classes depending on their topology within the membrane (Fig 1A). Type I proteins possess a single transmembrane domain (TMD) with a cytosolic C-terminus; these proteins also contain a cleavable ER targeting signal peptide at the N-terminus [6]. Type II proteins also contain a single TMD but display an opposite topology with a cytoplasmic N-terminus. The TMD of these proteins acts as a signal anchor that targets these proteins to the ER [7]. Type III proteins also have a single TMD and possess the same topology as type I proteins, but rather than containing a cleavable signal peptide their TMD acts as a reverse signal anchor [7]. Polytopic proteins possess multiple membrane-spanning domains, may or may not contain a cleavable signal peptide, and can possess either a luminal or cytoplasmic N- or C-terminus [7]. Finally, tail-anchored proteins have the TMD at the extreme C terminus and have their N terminus within the cytosol [8]. Examples of all types of transmembrane proteins are present in enveloped viruses.

**Figure 1.**
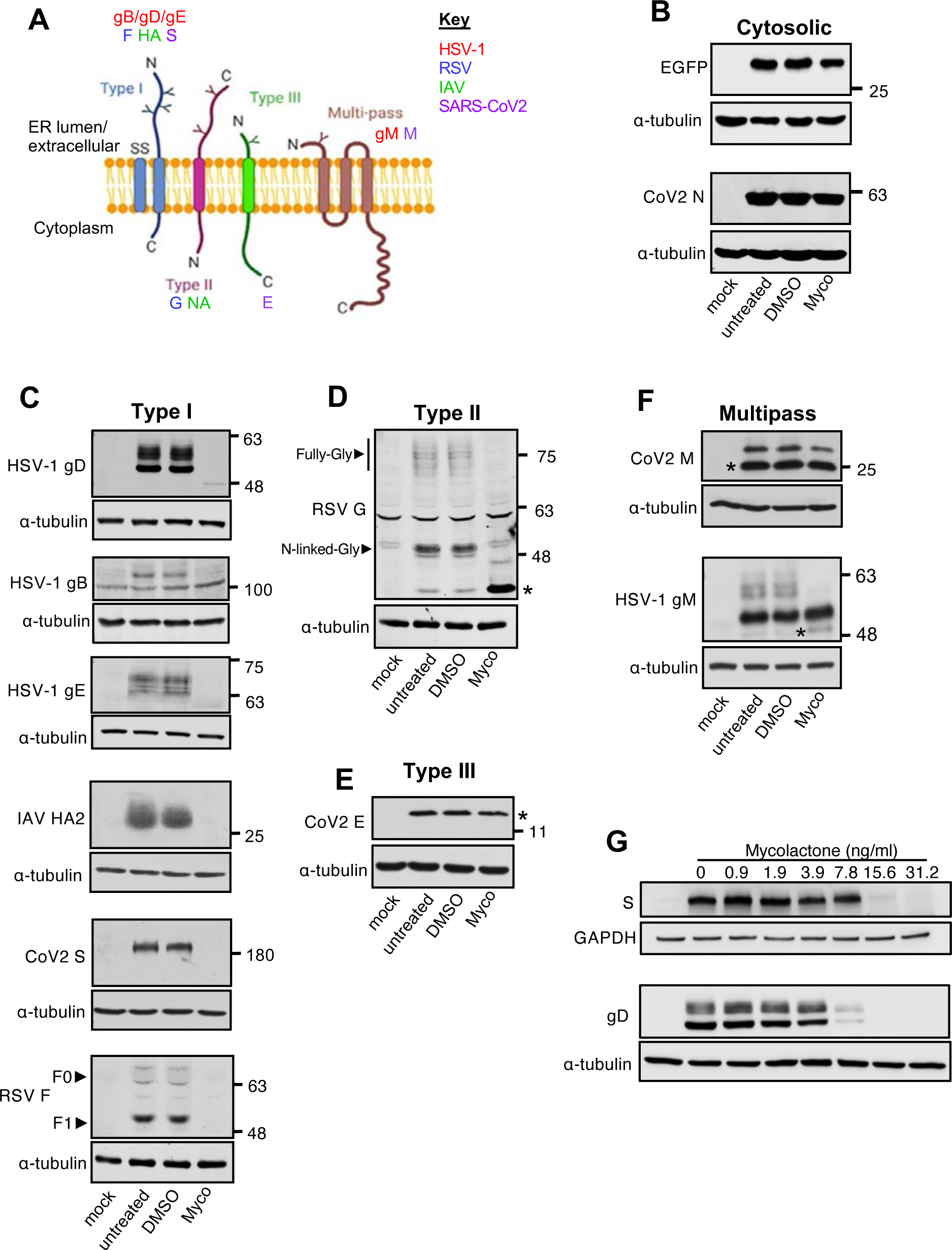
Mycolactone prevents biosynthesis of ectopically expressed virus envelope transmembrane proteins. (A) Cartoon illustrating topology of virus membrane proteins studied here. Type I membrane proteins with a cleaved signal peptide, external N terminus and internal C terminus. Type II proteins with a signal anchor transmembrane domain, internal N terminus and external C terminus. Type III membrane proteins with an external N terminus but no signal peptide/anchor and multipass, polytopic membrane proteins which cross the membrane multiple times. (B) to (F). HEK-293T cells were mock transfected or transiently transfected with plasmids expressing proteins of interest, and simultaneously treated with either 0.05% DMSO or 31.25ng/ml mycolactone (Myco). After 16 hours cells were harvested and analysed by SDS-PAGE and Western blotting for proteins as follows: (B) control proteins eGFP and SARS-CoV2 N; (C) type I membrane proteins HSV-1 gD, gB, gE, IAV HA, SARS-CoV2 S and RSV F; (D) type II membrane protein RSV G; (E) type III membrane protein SARS-CoV2 E and IAV HA; (F) multipass membrane proteins SARS-CoV-2 M and HSV1 gM. (G) Mycolactone dose response in HEK-293T cells transiently transfected with a SARS-CoV2 S or HSV1 gD encoding plasmid as described above. * indicates unglycosylated proteins.

Many transmembrane proteins are translocated into the ER membrane via the Sec61 translocon, a heterotrimeric complex that, together with its accessory proteins, forms a pore across the ER membrane allowing translocation of secretory proteins as well as type I and type II transmembrane proteins, and those polytopic proteins that have a signal peptide [9]. Other types of transmembrane proteins utilise other pathways, such as tail-anchored proteins that use the Get3/TRC40 pathway [8], and type III and polytopic proteins lacking signal peptides that use the EMC and PAT complexes respectively [10-12]. Regardless of the entry mechanism, translocation into the ER membrane is followed by trafficking through the secretory pathway which in the case of virus infection directs them to sites of virus envelopment, which vary across virus families but could be at the Golgi apparatus, the plasma membrane or the endocytic pathway [13]. Concomitant modification by glycosylation occurs in the ER and the Golgi apparatus [14, 15].

A potent inhibitor of the Sec61 translocon is an exotoxin from *Mycobacterium ulcerans* called mycolactone [16, 17]. Using cryo-electron microscopy, mycolactone has been shown to “wedge” into the so-called lateral gate of the Sec61 translocon. This is thought to stabilise the structure and prevent the signal sequence from binding, which is required for channel opening [18]. Mycolactone has been shown to inhibit the translocation of proteins that rely on Sec61 for their translocation; i.e. secreted proteins, and type I and type II transmembrane proteins, as well as signal peptide-bearing polytopic proteins [19, 20]. Inhibition of protein translocation means that these proteins are translated in the cytosol, where they are rapidly degraded by the proteasome [16] and/or removed by the autophagy machinery [21]. As a result, target protein production is ablated by mycolactone, leading to strong reductions in detection of newly expressed protein. Moreover, since many Golgi-resident enzymes required for second-order glycosylation are type II transmembrane proteins, exposed cells also rapidly lose the ability to undertake these modifications [22].

In this study we set out to determine if blockade of the Sec61 translocon by mycolactone could inhibit the expression of virus glycoproteins from several enveloped viruses that produce their envelope proteins from mRNAs transcribed using different strategies: the large DNA virus HSV1, which uses host cell machinery to transcribe mRNA from the double-stranded DNA genome in the nucleus; the negative-sense RNA virus RSV, which uses its own RNA-dependent RNA polymerase (RdRp) to transcribe its mRNA from the RNA genome in the cytoplasm; the segmented negative-sense RNA virus IAV, which uses its own RdRp to transcribe mRNAs from individual RNA genome segments in the nucleus; the positive-sense RNA virus SARS-CoV2 which transcribes its membrane protein mRNAs from the anti-sense genome in the cytoplasm; and the positive-sense RNA virus ZIKV which translates its positive-sense RNA genome into a single polyprotein on the ER membrane before cleavage into transmembrane and non-membrane proteins.

In line with its activity on cellular proteins, mycolactone inhibited the ectopic expression of all viral type I transmembrane proteins tested, while having no effect on type III or signal peptide-lacking polytopic proteins. *In vitro* translocation assays confirmed that lack of expression of type I proteins in cells was due to a failure of these proteins to translocate into the ER membrane. For the single type II protein tested (RSV G), Sec61-dependent translocation was blocked as expected, but unglycosylated protein was still detected. Mycolactone treatment of cells infected with these viruses also inhibited expression of their type I membrane proteins, and caused the mislocalisation of unglycosylated RSV G. The unaltered expression of type III, polytopic or proteins that were translated in the cytosol indicated that the different biogenesis pathways of these proteins are insensitive to mycolactone. However, although mycolactone treatment was also shown to inhibit downstream virus production, the window for mycolactone sensitivity during virus infection was unexpectedly restricted to very early times in life cycles of some of these viruses, in particular for IAV, with efficient virus propagation occurring before virus transmembrane protein synthesis was detectable. This reveals that the morphogenesis of some enveloped viruses requires limited amounts of envelope proteins and that the large amounts ultimately produced during virus infection are in vast excess in the infected cell and likely have additional roles.

## Results

### Impact of mycolactone on host cell viability

We have employed a broad range of primary and established human and animal host cell types in this study: African green monkey kidney lines BS-C-1 and Vero, canine kidney MDCK cells, the human epithelial line HeLa and its derivative HEp2, human embryonic kidney HEK-293T cells, human colonic epithelial Caco-2 cells, human keratinocytes HaCaT cells and primary human foreskin fibroblasts, HFFF. To establish the dose required to completely inhibit Sec61 in each of these cell types, cytotoxicity was first determined using established protocols [23] after five days of exposure (Fig S1A to S1I). In all cell types, mycolactone reduced cell viability at low nanomolar concentrations, but the minimum inhibitory concentration (MIC) values, asterisked in Fig S1, varied from 3.91ng/ml (HFFF, Caco-2 and HEp2) to 31.25ng/ml (BS-C-1 and Vero). The IC_50_ values followed a similar pattern ranging from 0.45ng/ml (HFFF) to 4.31ng/ml (Vero) (Fig S1J). Short term exposure to mycolactone for 24 hours, at or above the MIC had a limited impact on viability in most cell types (Fig S2). Hela, MDCK and HaCaT cells were more sensitive to mycolactone with viability dropping to around 80%, while HFFF cells proved to be the most sensitive with 31.25ng/ml mycolactone sufficient to cause a 50% drop in viability. This limited effect of short exposure to mycolactone in most cell types gave us confidence that changes in viral protein production and replication over this timeframe could be attributed to the direct effects of mycolactone rather than secondary effects of host cell death.

### Ectopic expression of type I transmembrane proteins from a range of enveloped human viruses is blocked by mycolactone

The four types of transmembrane protein are all represented by membrane proteins expressed by the viruses used in this study (Fig 1A), but the size and complexity of these proteins varies considerably (Fig S3). For example, the F protein of RSV is synthesised as an F_0_ precursor which is cleaved into an F_1_ fragment by a furin-like protease in the Golgi, while HSV1 glycoproteins do not undergo further processing after translocation. The ability of mycolactone to block expression of these proteins when expressed in isolation was tested by transiently transfecting HEK-293T cells with plasmids expressing proteins that do not use Sec61 in their biogenesis as they are translated in the cytosol (GFP, SARS-CoV2 N), or the membrane proteins indicated in Fig 1A in the presence of DMSO or mycolactone.

Western blotting of transfected cell lysates indicated that, as predicted, neither of the cytosolically expressed proteins was affected by mycolactone treatment (Fig 1B). However, expression of all type I membrane proteins was abrogated by mycolactone (Fig 1C). Mycolactone did not alter the overall level of RSV G expression, the only type II membrane protein studied here by ectopic expression but dramatically affected the relative migration of expressed protein (Fig 1D). RSV G protein is known to have 30 to 40 O-linked glycans in addition to 4 to 5 N-linked glycosylation sites [24], and these account for the large molecular weight of the mature fully-glycosylated form of the protein (∼ 80 kDa), while the partially-glycosylated variant at ∼ 48 kDa likely represents the N-linked glycosylated protein prior to addition of O-linked glycosylation. By contrast, the protein detected in the presence of mycolactone was the correct size for the non-glycosylated form of RSV G (∼33 kDa). Taken together with previous data, it seems likely that RSV G did not access the secretory pathway and instead was translated in the cytosol in the presence of mycolactone but is unusually resistant to proteasomal degradation. Intriguingly, the expression of the SARS-CoV2 E protein was not reduced in the presence of mycolactone (Fig 1E), confirming its topology as a type III membrane protein [25]. Likewise, the polytopic signal peptide-lacking multipass SARS-CoV-2 M and HSV1 gM proteins were still expressed in mycolactone-exposed cells, although in the case of gM, the fully glycosylated protein was absent in the presence of mycolactone (Fig 1F) likely due to the known effect of mycolactone on the glycosylation machinery [22]. The inhibition of HSV1 gD and SARS-CoV2 S expression were confirmed to be dose dependent (Fig 1G).

To determine if mycolactone affects the localisation of those membrane proteins that were still expressed after treatment, transfected HeLa cells were fixed and stained for immunofluorescence (Fig 2). In line with results from Western blotting, mycolactone treatment had no effect on the diffuse cytoplasmic localisation of the SARS-CoV-2 N protein (Fig 2, N). However, the type I membrane protein HSV1 gD, which was localised to the secretory pathway in the absence of mycolactone, was barely detectable in its presence (Fig 2, gD). In the case of the type III SARS-CoV2 E protein, mycolactone had no effect on its expression (Fig 1E) or localisation (Fig 2, E), where it was detected in distinctive domains within the cytoplasm. Although the Golgi apparatus is fragmented in cells expressing E, these domains do not represent fragmented Golgi (Fig S4, E) but are likely to be the ER-Golgi intermediate compartment as found for ectopic expression of SARS-CoV E protein [26]. Likewise, the localisation of the polytypic multipass proteins SARS-CoV2 M and HSV1 gM to the secretory pathway was unaffected by mycolactone treatment (Fig 2, M and gM), with M concentrated in the Golgi (Fig S4, M) and gM distributed through the secretory pathway (Fig 2, gM). This localisation data confirms the high sensitivity of type I membrane protein biogenesis to mycolactone.

**Figure 2.**
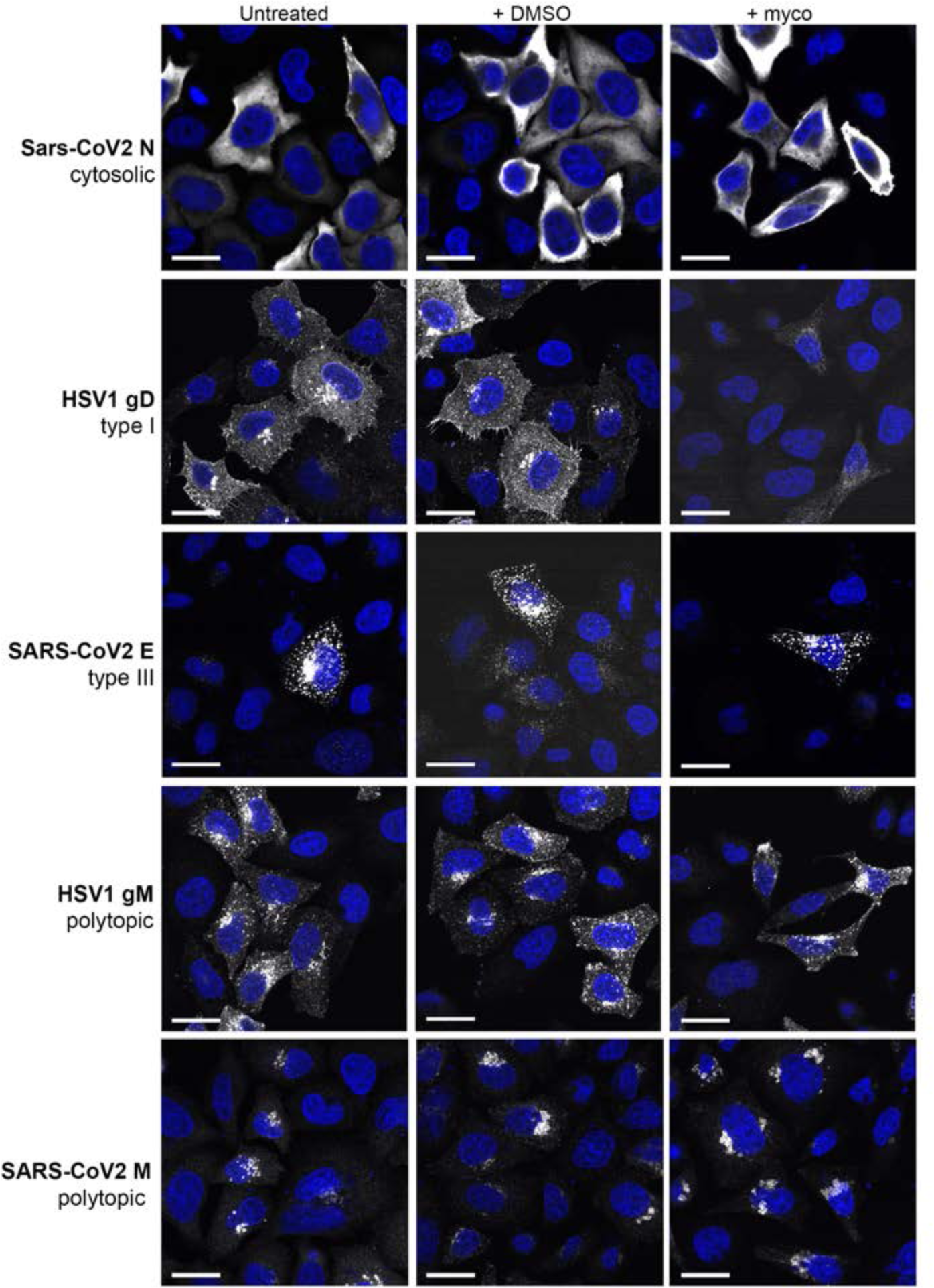
Subcellular localisation of ectopically expressed virus-encoded transmembrane proteins in the presence of mycolactone. HeLa cells were transiently transfected with plasmids expressing HSV1 gD (type I), SARS-CoV2 E (type III), HSV1 gM, SARS-CoV2 M (both multipass) or SARS-CoV2 N (control), and left untreated or simultaneously treated with either 0.05% DMSO or 31.25ng/ml mycolactone (Myco). After 16 hours cells were fixed and stained for the indicated proteins (white) and the nuclei stained with DAPI (blue).

### Mycolactone inhibits Sec61-directed translocation of viral type I and II membrane proteins *in vitro*

To directly investigate the effect of mycolactone on the translocation of viral glycoproteins, we carried out *in vitro* translation/translocation assays using canine pancreatic microsomal membrane preparations (CPMM) (Fig 3). In the absence of CPMM, the type I transmembrane HSV1 gD was synthesized as a single polypeptide, while in the presence of membranes, translocation of the nascent peptide chain allowed glycosylation and a higher molecular weight band, which was unchanged when the membranes were preincubated with DMSO (Fig 3A). Preincubation of the CPMM with mycolactone on the other hand led to a complete block in glycosylation with only the non-glycosylated protein detectable (Fig 3A). This confirms that mycolactone blocks the translocation of nascent gD into the ER via Sec61, and in transfected cells this likely results in degradation of the non-translocated gD by the proteasome.

**Figure 3.**
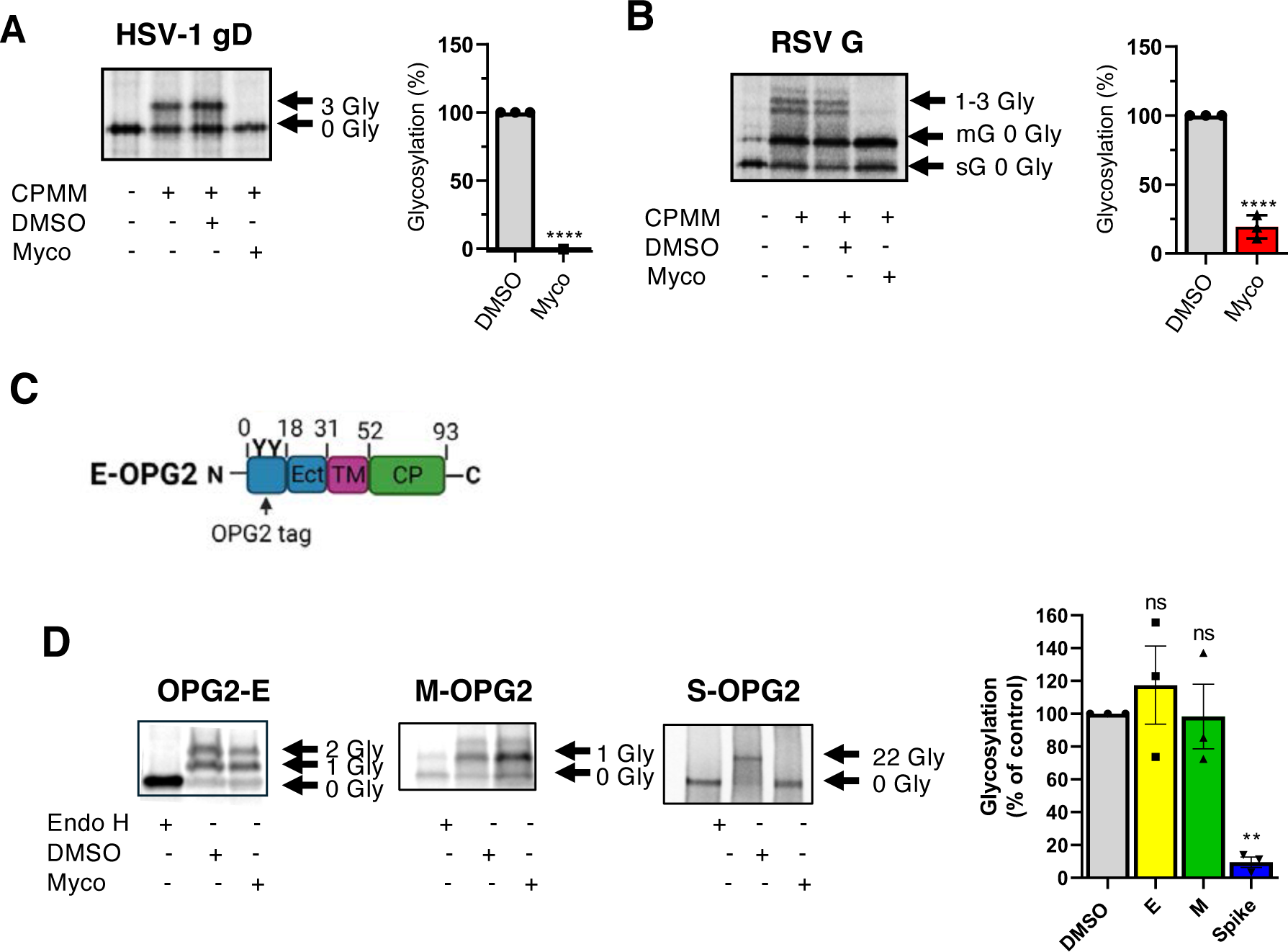
Inhibition of viral envelope glycoprotein expression by mycolactone is due to the blockade of translocation. *In vitro* translocation assays of ^35^S labelled HSV-1 gD (A) or RSV G (B) translated from mRNA transcripts in the presence or absence of CPMM incubated with DMSO or 200ng/ml mycolactone. Products were subjected to SDS-PAGE and imaged with a phosphorimager and representative images are shown. Quantitation is calculated using the ratio of N-glycosylated to non-glycosylated protein relative to the untreated CPMM control. Data is presented as mean±SEM, n=3 independent repeats. Statistical significance comparing the mycolactone treatment to the untreated control was assessed using a two-tailed unpaired *t* test. Statistical significance: *p* <0.0001; ****. (C) Cartoon of OPG2-tagged SARS-CoV2 E protein. (D) In vitro translocation assays using the pcDNA5-OPG2-E, pcDNA5-M-OPG2 or pcDNA5-S-OPG2 plasmids used directly within a TnT® coupled transcription/translation system. Assays were carried out in the presence of CPMM exposed to DMSO or 200ng/ml mycolactone. Membrane fractions were left untreated or subjected to EndoH digestion before SDS-PAGE separation and phosphoimager analysis. Glycosylation ratios calculated as described above. Data is presented as mean±SEM, n=3 independent repeats. Statistical significance comparing the mycolactone treatment to the untreated control was assessed using a one-way ANOVA. Statistical significance: *p* <0.01; **.

Since mycolactone blocked the glycosylation of the type II protein RSV G without affecting overall abundance during ectopic expression, we used this assay as the gold standard to determine whether translocation of this substrate is blocked by mycolactone. This analysis was complicated by the presence of an alternative translation start site in the mRNA resulting in an additional a smaller soluble form that has no transmembrane region (sG) as well as the higher molecular weight type II protein(mG) (Fig S3B). In the presence of CPMM, three glycosylated protein bands appeared which corresponded to the three potential N-linked glycosylation sites in the ectodomain (Fig 3B), and all of these were lost when mycolactone was present. Therefore, translocation of RSV G was inhibited by mycolactone, supporting our contention that the non-glycosylated G observed in mycolactone-treated cells (Fig 2D) has likely not accessed the ER.

In the case of HSV gD and RSV G assays above, endogenous glycosylation sites in the protein were utilised to determine protein translocation. To analyse translocation of the SARS-CoV2 proteins, we utilised the OPG-2 tagged proteins described elsewhere [25] which add an N-glycosylation site that allows detection of translocation in proteins such as the SARS-Cov-2 E protein that lack endogenous glycosylation signals (Fig 3C). In these assays, membranes were isolated by centrifugation to allow concentration of the translocated proteins, and the presence of glycosylated proteins confirmed by the shift in molecular weight following EndoH digestion (Fig 3D). Similar levels of E and M protein glycosylation were observed in both the presence and absence of mycolactone, indicating that mycolactone was unable to inhibit their translocation, as predicted by their topology and in line with cell transfection data. By contrast, mycolactone completely blocked the glycosylation of the type I S protein.

A similar effect on the glycosylation of S was seen when membranes were incubated with a structurally dissimilar Sec61 inhibitor, Coibamide A, confirming that the effect is due to translocation inhibition (Fig 4A). To further confirm the Sec61 dependence of the inhibition of the S protein synthesis, mycolactone, Coibamide A and a second unrelated Sec61 inhibitor, Ipom F, which has previously been shown to block S protein translocation in *vitro* [25], were included in HEK293-T transfection assays (Fig 4B). All three compounds showed a similar level of inhibition of S protein expression in a cellular context. Finally, to confirm the Sec61-dependence of the inhibitory effect of mycolactone on S protein production, transfections were carried out in HEK293T cells expressing a Flag-tagged Sec61 mutant, D60G, that is unable to bind mycolactone and is therefore resistant to inhibition [18]. No inhibition of transient S protein expression was observed in cells containing D60G (Fig 4C). Taken together, these data confirm that the effect of mycolactone on virus envelope protein expression is due to its action at the Sec61 translocon.

**Figure 4.**
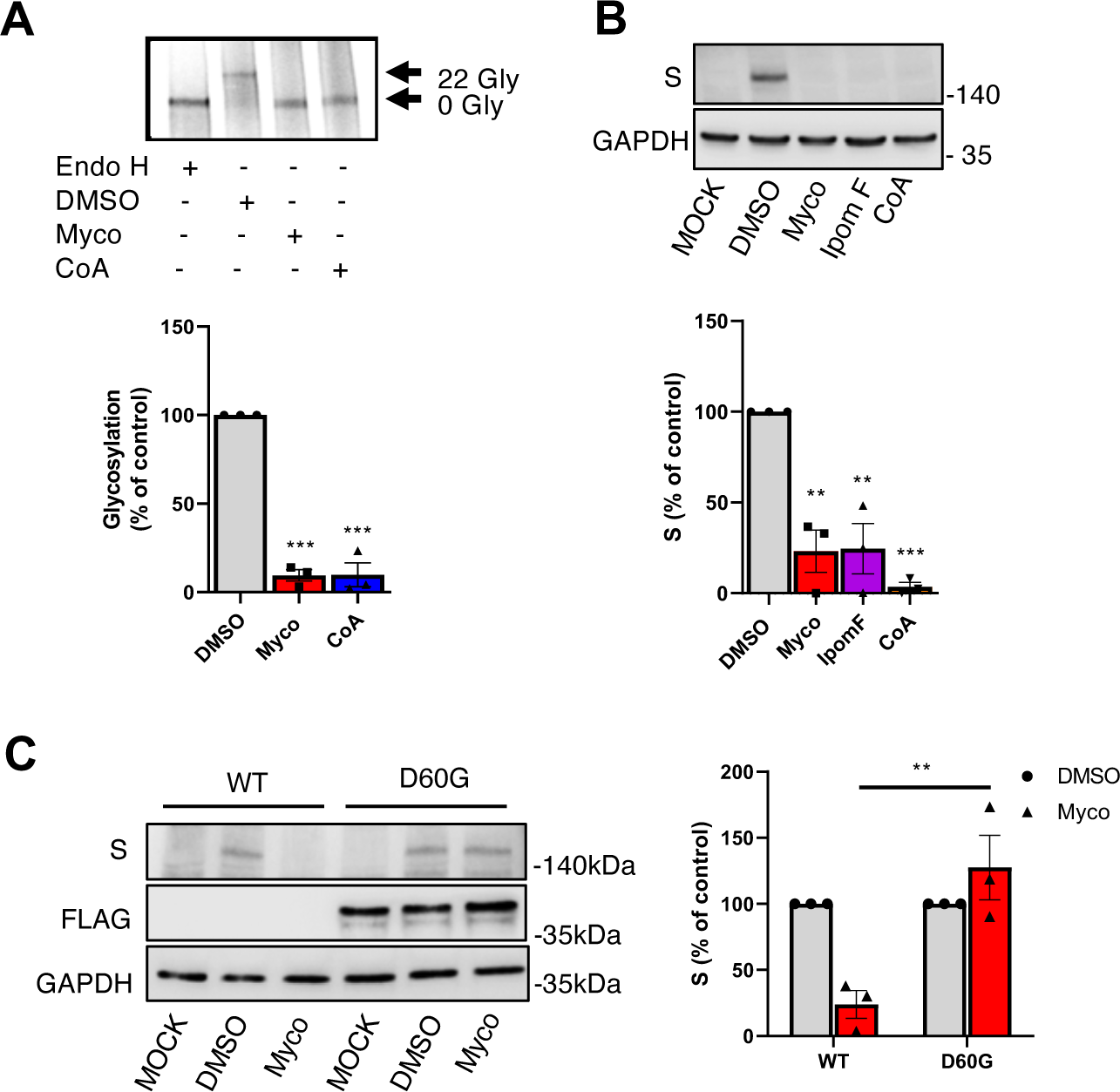
The inhibition of virus envelope glycoprotein expression by mycolactone is a consequence of Sec61 inhibition. (A) In vitro translocation assays of pcDNA5-S-OPG2 using CPMM preincubated with DMSO, 200ng/ml mycolactone (Myco) or 1µM Coibamide A (CoA). Quantitation was calculated using the ratio of N-glycosylated to non-glycosylated protein relative to the untreated CPMM control. Data is presented as mean±SEM, n=3 independent repeats. Statistical significance comparing the mycolactone treatment to the untreated control was assessed using a one-way ANOVA. Statistical significance: *p* <0.001; ***. (B) Western blots of SARS-CoV-2 S protein expression in transiently transfected Hek-293T cells exposed to 31.25ng/ml mycolactone (myco), 250nM Ipomoeassin F (Ipom F) or 500nM Coibamide A (CoA) for 16hr. Quantitative data is normalised according to GAPDH and presented as percentage of DMSO control, mean±SEM, n=3 independent repeats. Statistical significance was assessed using a one-way ANOVA. Statistical significance: *p*<0.01; ** *p* <0.001; ***. (C) Western blots of SARS-CoV-2 S protein expression in transiently transfected WT Hek-293T and cells expressing the mycolactone resistant Flag-tagged SEC61A1 D60G mutant exposed DMSO or 31.25ng/ml mycolactone (Myco). Quantitative data is normalised according to GAPDH and presented as percentage of DMSO control, mean±SEM, n=3 independent repeats. Statistical significance was assessed using a two-tailed unpaired *t* test. Statistical significance: *p* <0.01; **.

### Host-directed targeting of Sec61 translocation during virus infection by enveloped viruses

Given the profound effect of mycolactone on the expression of type I membrane proteins, we predicted that mycolactone would abrogate virus production for any enveloped virus that relied on type I membrane proteins for their envelopment and/or entry into cells. To first test if mycolactone had the capacity to block the production of virus envelope proteins when expressed from their own genomes, the appropriate cells were pretreated with DMSO or mycolactone for 30 mins or left untreated, then infected with HSV1, RSV, IAV or SARS-CoV2 (Fig 5A to 5D). In all cases, the virus membrane proteins exhibited the same relative sensitivity to mycolactone during virus infection as shown above during ectopic expression, with type I membrane protein expression blocked (HSV1 gB, gD, gE; RSV F; IAV HA; SARS-CoV2 S). Moreover, our results with IAV proteins are in agreement with a previous study where the cell surface levels of ectopically expressed virus membrane proteins were measured in the presence of mycolactone [19]. Intriguingly, HSV1 gB persisted at a reduced expression level in its non-glycosylated form in the presence of mycolactone during infection, despite its expression being fully blocked by the action of mycolactone when expressed ectopically (compare gB in Fig 5A with gB in Fig 1C). While the type II IAV NA protein expression was blocked by mycolactone, RSV G was resistant and found in an unglycosylated form, in line with its ectopic expression above (Fig 1). The polytopic (HSV1 gM; SARS-CoV2 M) proteins were resistant to the action of mycolactone, while the efficient expression of non-membrane (RSV N; IAV NP; SARS-CoV2 N) proteins indicated that entry and gene expression of these viruses were unaffected by mycolactone treatment. Importantly, mycolactone pretreatment blocked virus production of all these viruses, as expected from the absence of type I membrane protein expression (Fig 5E to 5H).

**Figure 5.**
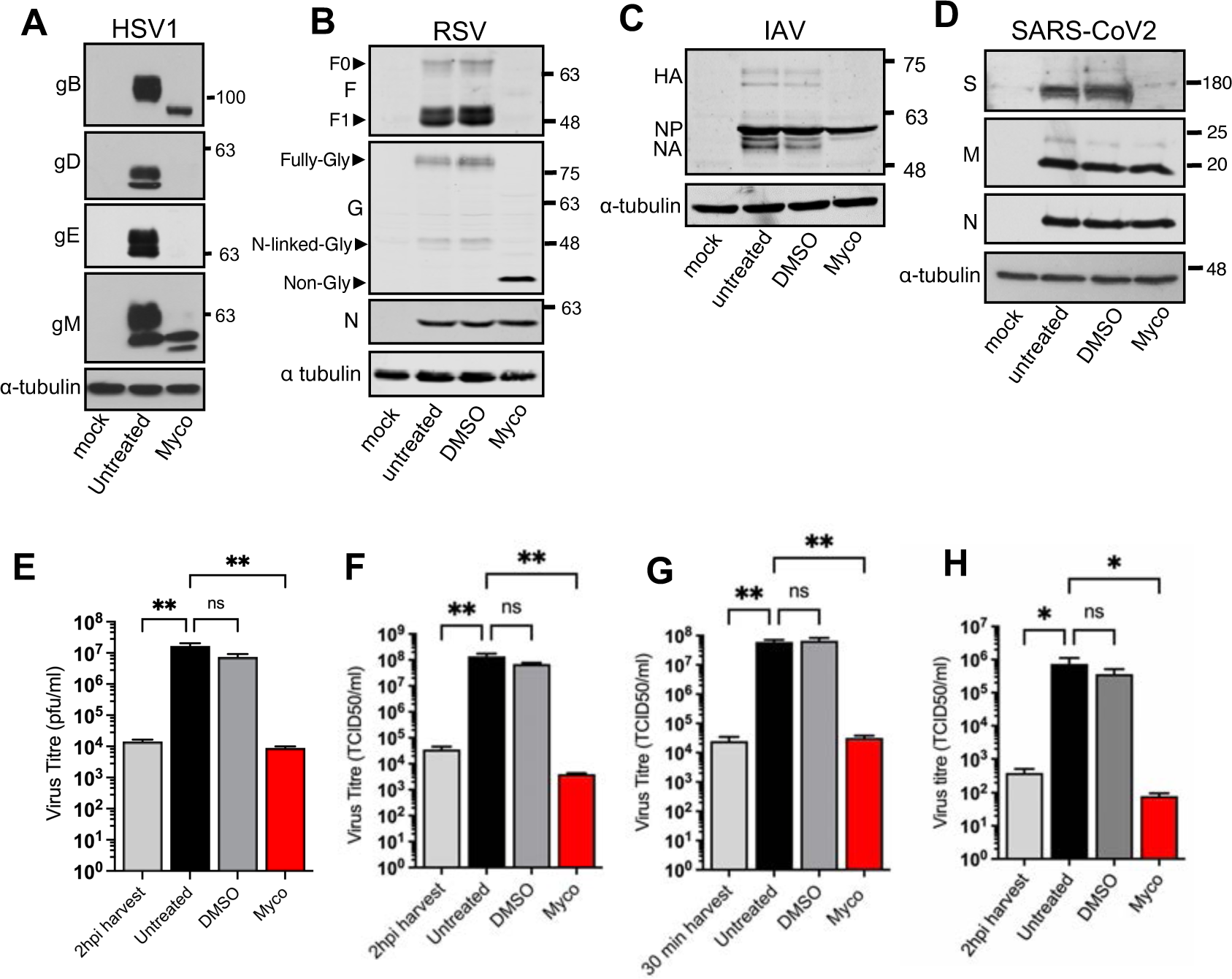
Mycolactone blocks virus envelope protein synthesis and inhibits production of infectious virus. (A) to (D) HFFF, Hep2, MDCK or Caco-2 cells were pretreated for 30 minutes with either 0.05% DMSO or 31.25ng/ml of mycolactone, before being infected with HSV1, RSV, IAV (all at MOI 2) or SARS-CoV2 (MOI 0.2) respectively. Mycolactone and DMSO were additionally present during the inoculation period and throughout the infection. Total cell lysates were harvested at appropriate times after infection (HSV1,16 hpi; RSV, 22 hpi; IAV, 10 hpi; SARS-CoV2, 18 hpi) and subjected to SDS-PAGE and Western blot for virus proteins as indicated, using α-tubulin as loading control. Differentially glycosylated forms of RSV F and G are indicated with arrowheads in (B). (E) to (H). Infections were carried out as for (A) to (D) and total virus was harvested and subjected to titration by plaque assay (HSV1) or TCID50 titration (RSV, IAV and SARS-CoV2). Data is presented as mean±SEM, n=3 biological repeats. In all cases, statistical significance was assessed using one-way ANOVA and Dunnett’s multiple comparisons test. Statistical significance: **, *p* <0.01; *\*, p* < 0.05; ns, non-significant.

Using HSV1, we next conducted a detailed investigation of the effect of mycolactone when applied to infected cells subsequent to virus infection. HFFF cells were infected with HSV1 at a multiplicity of 2, treated after 3 hours with increasing doses of mycolactone, and total virus harvested after 16 hours for titration on Vero cells. Mycolactone had little effect on virus production at lower doses, but addition of > 30 ng/ml resulted in a three to four-log reduction in virus titre (Fig 6A). Infection of HaCaT keratinocyte cells treated with the same concentration of mycolactone also resulted in close to a five-log reduction in virus titre, with infection of a range of other cell types showing a similar inhibition (Fig S5). We next wished to determine how long HSV1 remained sensitive to mycolactone treatment during the virus replication cycle. HSV1 replicates relatively rapidly in HaCaT cells [27], with the production of progeny virus increasing rapidly up to 16 hours (Fig 6B). Addition of mycolactone at different times of infection confirmed that before six hours, mycolactone blocked new virus production effectively but after six hours, virus production increased in line with the one-step growth curve (Fig 6C), indicating that once progeny virions had been assembled, they were no longer sensitive to the action of mycolactone. However, in the corresponding Western blots of cell lysates for this experiment, mature glycosylated gE and gM were only detectable when mycolactone was added from 8 hours onwards, while type I membrane proteins gB and gD were not detected in their mature, glycosylated forms at any time of mycolactone addition before 12 hours, suggesting that the translocation and trafficking through the secretory pathway of the majority of glycoproteins synthesized during virus infection occurred late in infection (Fig 6D). This contrasted with the two non-membrane virus structural proteins that were tested, VP22 and VP16, both of which were detected at high levels even when mycolactone was added as early as two hours. These data suggest that some essential glycoproteins are undetectable in their mature, secretory pathway forms at times when there is already active virus production, and that, as we have suggested previously [28], HSV1 glycoproteins appear to be made in significant excess over the levels required for virus morphogenesis.

**Figure 6.**
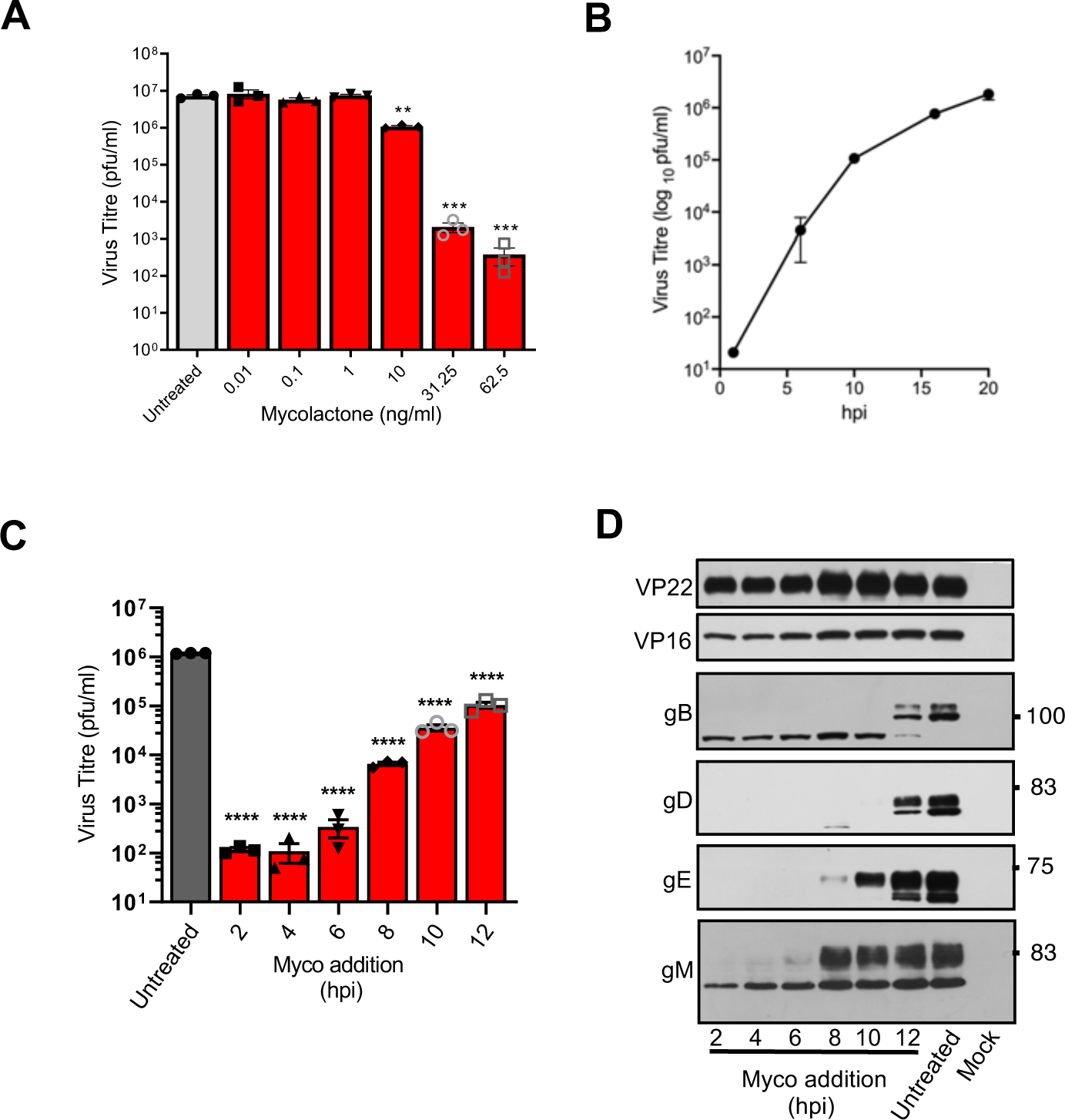
Inhibition of HSV1 production by mycolactone. (A) HFFF cells were infected with HSV1 at a multiplicity of 2 and treated with increasing concentrations of mycolactone at 3 hpi and harvested at 20h and titrated on Vero cells. (B) One-step growth curve of HSV1 in HaCaT cells. (C) & (D) HaCaT cells were infected with HSV-1 at a multiplicity of 2, and at the indicated times were treated with mycolactone (Myco). At 14 hours post infection cells were harvested for virus titration (C) or Western blotting (D). Data is presented as mean±SEM, n=3 biological repeats. Statistical significance comparing the mycolactone treatment to the untreated control was assessed using one-way and Dunnett’s multiple comparisons test. Statistical significance: p <0.0001;****.

### Mycolactone treatment of cells infected with negative-stranded RNA viruses

HSV1 is a highly complex virus that encodes over 80 proteins including at least ten envelope proteins, with scope for redundancy in the system. To determine the efficacy of mycolactone on additional enveloped viruses, we next tested its effect on RSV, a small, enveloped virus with a negative sense RNA genome that replicates in the cytoplasm and expresses just one essential glycoprotein, the type I F glycoprotein. To establish RSV growth kinetics in our system, a one-step growth curve of RSV strain A2 was carried out in HEp2 cells infected at an MOI of 2 (Fig 7A). RSV replication occurred more slowly than HSV1, with virus production increasing rapidly between 12 and 24 hours before peaking at 48 hours. Western blotting of infected cell extracts indicated that glycoproteins F and G and the cytosolic N protein were all barely detectable at 9 (Fig 6B) with virus protein synthesis peaking between 12 and 24 hpi, in line with the one-step growth curve.

**Figure 7.**
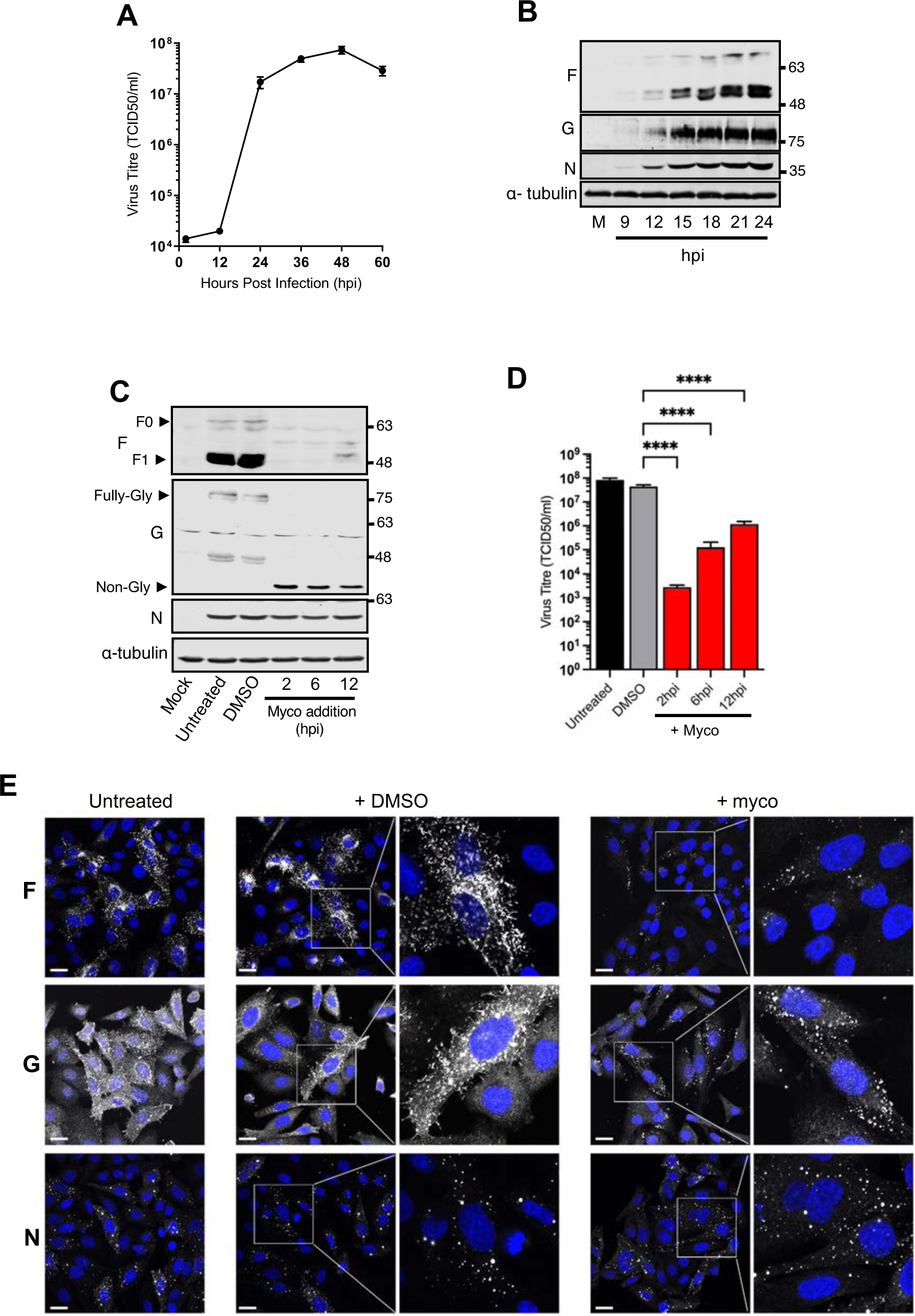
Outcome of RSV infection after treatment with mycolactone. (A) HEp2 cells were infected with the RSV A2 strain at MOI 2. At the indicated times total virus was harvested for TCID50 titration on HEp2 cell monolayers. Data is presented as mean±SEM, n=3 biological repeats. (B) Total cell lysates from HEp2 cells infected as above, collected at the indicated times, were analysed by SDS-PAGE and Western blotting for RSV proteins F, G and N; α-tubulin was used as a loading control. (C) & (D) RSV infected Hep2 cells were treated with mycolactone at the indicated times and harvested at 24 hours for Western blotting (C) or TCID50 titration on Hep2 cells (D). Data is presented as mean±SEM, n=3 biological repeats. Statistical significance comparing the mycolactone treatment to the DMSO control was assessed using one-way ANOVA and Dunnett’s multiple comparisons test. Statistical significance: p <0.01; **. (E) HEp2 cells seeded onto glass coverslips were infected with RSV at a MOI 2, fixed and stained for the indicated proteins (white) and the nuclei were stained with DAPI (blue). Scale bar = 20 μm.

To test how long RSV infected cells remain sensitive to Sec61 blockade, infected cells treated with mycolactone at various times up to 12 hours were tested for protein synthesis and virus production. As late as 12 hours, F protein was undetectable and G was in its non-glycosylated form, but N was synthesized at normal levels (Fig 7C). Intriguingly, unlike the situation in HSV1 infection, addition of mycolactone at 12 hours (before virus production had begun and F protein was undetectable) reduced the overall yield of RSV by only 10 to 100 fold (Fig 7D, 12 h), indicating that sufficient but undetectable levels of virus envelope proteins had entered the secretory pathway and reached the plasma membrane to allow capsid envelopment to occur later in infection.

To determine the effect of mycolactone on RSV structural protein localisation, and in particular to better understand that of the non-glycosylated G protein, the cellular location of the viral proteins was examined by immunofluorescence at 14 hpi (Fig 7E). The F protein was localised in both the secretory pathway in its characteristic filaments at the plasma membrane (PM) in control and DMSO treated cells but was barely detectable in cells treated with mycolactone at 6 hours. By contrast, in the absence or presence of mycolactone, the N protein was present in its well-described cytoplasmic inclusion bodies which are the sites of viral RNA synthesis [29], confirming that blockade of the Sec61 translocon had no effect on expression of N or formation of inclusion bodies. The G protein was detected in punctate cytoplasmic structures that were different to inclusion bodies (Fig S7), and at the PM in control and DMSO treated cells. However, although G puncta were still formed in the presence of mycolactone, G did not localise to the plasma membrane, confirming that its trafficking through the secretory pathway and subsequent virus budding at the PM had been blocked. We do not yet understand the identity of these G-specific puncta, but since we showed that translocation into the ER of RSV G is blocked by mycolactone, and that expressed protein remains unglycosylated, it seems likely that they are outside the secretory machinery of the cells.

Unlike RSV and HSV1, IAV replicates rapidly in culture with virus production starting around 4 hours and peaking by 12 hours (Fig 8A). Nonetheless, protein production was barely detectable at 4 hours, but by 6 hours, significant amounts of virus proteins NP and M1 and to a lesser extent the type I membrane protein HA and the type II membrane protein NA were present in infected cells (Fig 8B), suggesting there was little lag time between protein synthesis and virus morphogenesis in IAV infected cells. As was the case with pretreated cells, mycolactone blocked the expression of HA but not the non-membrane proteins N or M1 (Fig 7C), confirming its specificity for the IAV transmembrane protein. The expression of the type II membrane protein NA was also blocked, in line with previous work showing the sensitivity of type II proteins to mycolactone [19, 20]. Strikingly, addition of mycolactone after infection had no effect on IAV production (Fig 7D). Moreover, addition of mycolactone as early as 30 mins after infection had little effect on IAV production (Fig 7E) suggesting that the virus had already produced sufficient membrane proteins which had entered the secretory pathway within this short time to facilitate virus budding and release at the plasma membrane in high quantities. Taken together, these data imply that IAV synthesizes sufficient glycoproteins within these first 30 mins to provide enough membrane protein for many assembled capsids to bud into later in infection.

**Figure 8.**
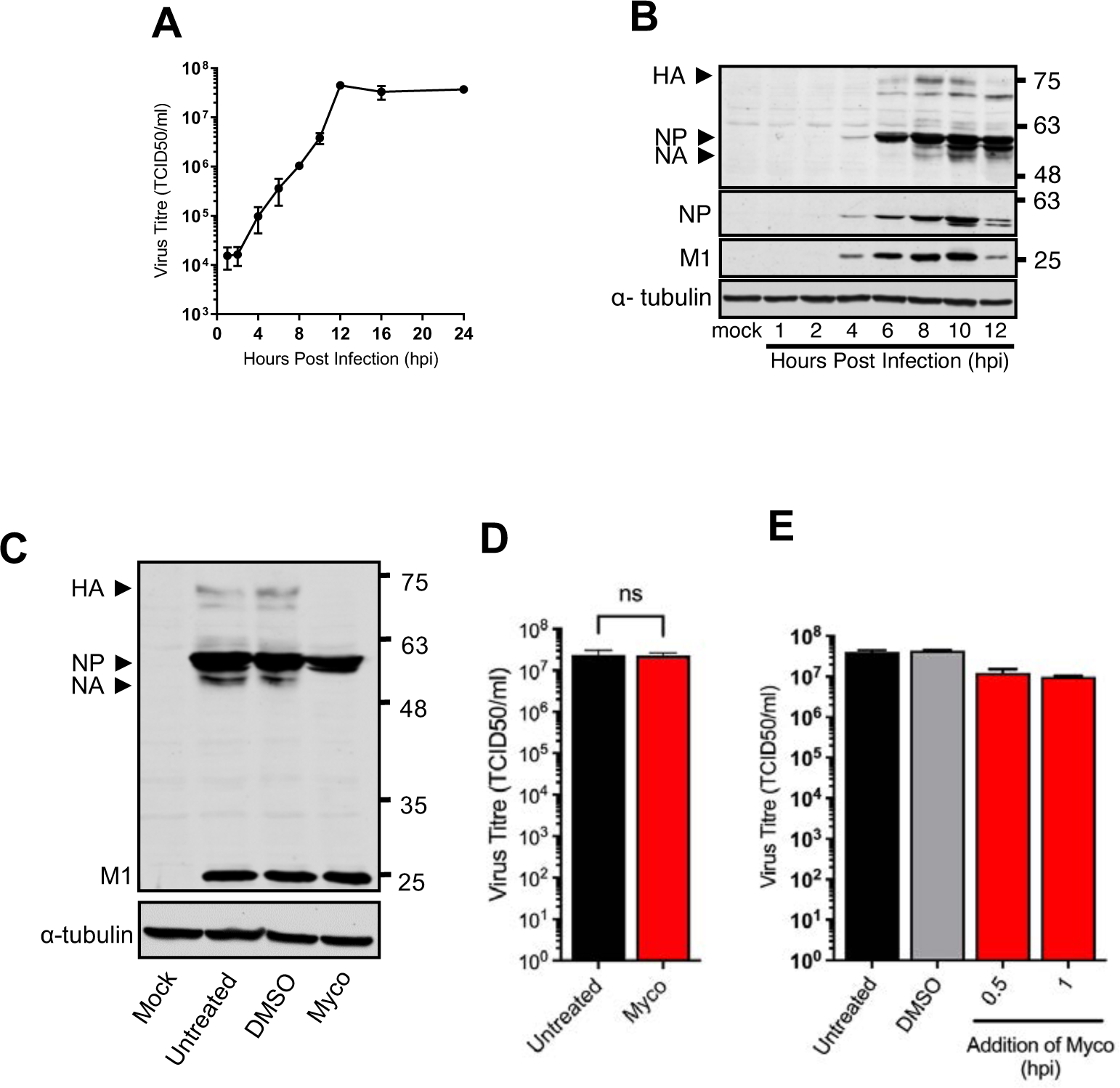
Production of IAV is relatively resistant to mycolactone. (A) MDCK cells were infected with IAV at MOI 2. At the indicated times total virus was harvested for TCID50 titration on MDCK cell monolayers. Data is presented as mean±SEM, n=3 biological repeats. (B) Total cell lysates from MDCK cells infected as above were harvested at indicated times, subjected to Western blot analysis and probed for the proteins indicated using the appropriate antibodies. (C), (D) & (E) MDCK cells were infected with IAV as described above, mycolactone was added at 31.25ng/ml at 2 hpi (D) or 0.5 and 1 hpi (E), with 0.05% DMSO added at 0.5 hpi used as a control. Total virus was harvested at 10 hpi for TCID50 titration on MDCK cells. Data is presented as mean±SEM, n=3 biological repeats.

### Mycolactone treatment of cells infected with a positive-stranded polyprotein-producing RNA virus

Positive-stranded RNA viruses such as ZIKV hijack ER membranes for their replication. Their RNA genomes are translated immediately upon infection into a large single polypeptide that is co- and post-translationally cleaved by host and viral proteases to produce individual viral proteins, six of which are integral membrane proteins including the membrane (M) and the envelope (E) structural proteins (Fig S3E). ZIKV replicates relatively slowly in culture with virus production and detectable protein synthesis of the cytoplasmic proteins C, NS3 and NS5, beginning after 12 hours (Fig 9A & 9B). Unlike the other viruses studied above, mycolactone treatment of infected cells early in infection (2 hours) fully blocked production of these cytoplasmic proteins, likely reflecting the requirement for the Sec61 translocon in the correct translation and orientation of the polyprotein in the ER membrane (Fig 9C). Consequently, virus production was also blocked by mycolactone treatment at this time (Fig 9D). Nonetheless, similar to RSV and IAV, delaying the addition of mycolactone to 12 hours, before virus production or detectable protein synthesis had begun, also rescued a significant proportion of infectious progeny (Fig 9D).

**Figure 9.**
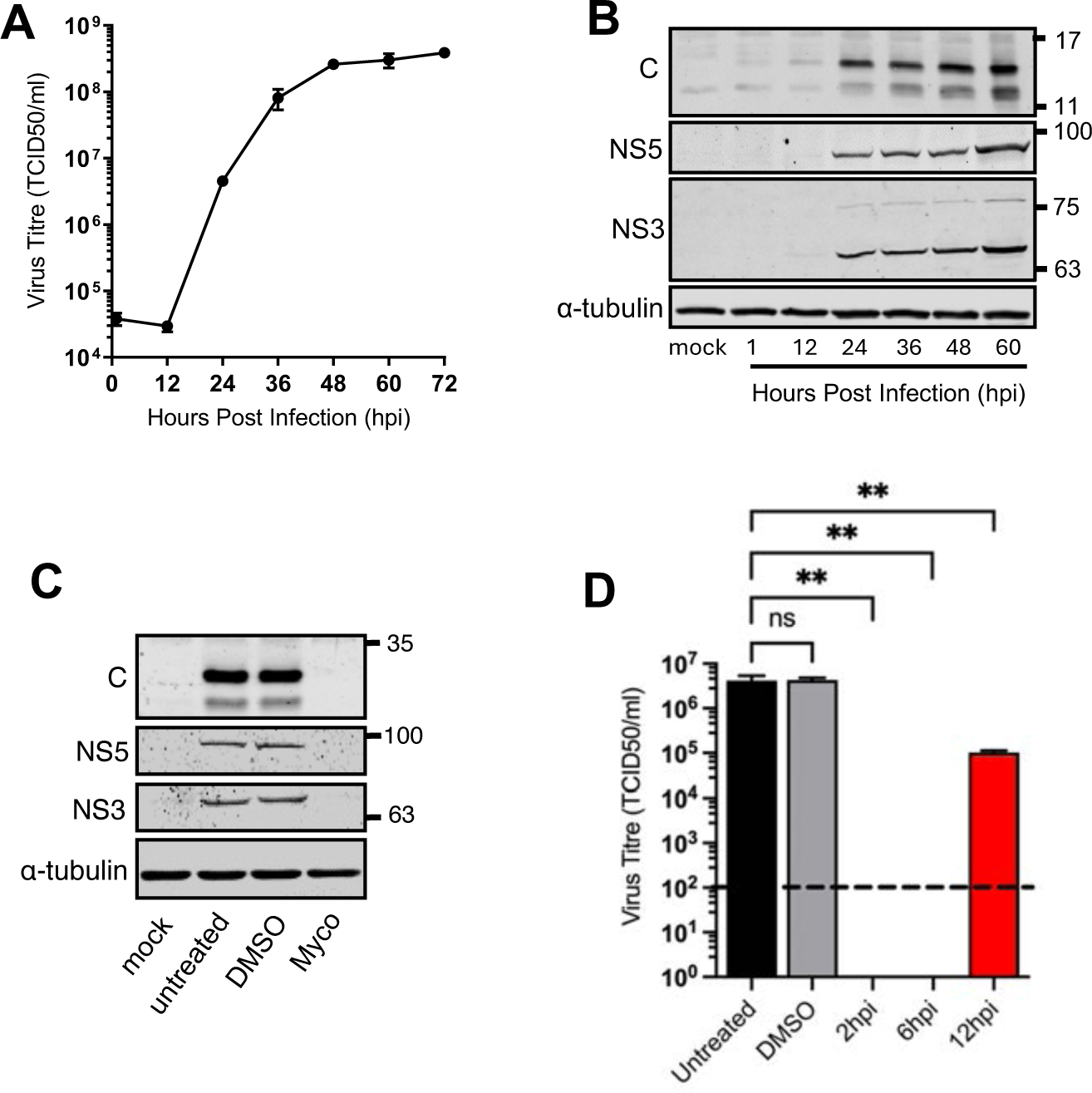
Mycolactone blocks expression of the ZIKV polyprotein. (A) Vero cells were infected with ZIKV at MOI 2. At the indicated times total virus was harvested and TCID50 titration was conducted on Vero cell monolayers to determine virus titres. Data is presented as mean±SEM, n=3 biological repeats. (B) Total cell lysates were subjected to SDS PAGE and Western blot analysis for the proteins indicated using α-tubulin as a loading control. (C) Vero cell monolayers were infected with ZIKV at MOI 2 and treated with 0.05% DMSO or 31.25ng/ml of mycolactone at 2 hpi. At 24 hours total cell lysates were harvested and subjected to SDS-PAGE and Western blot analysis for the proteins indicated. (D) Vero cell monolayers were infected with the ZIKV at MOI 2, and were either treated with 0.05% DMSO at 2 hpi or with 31.25ng/ml of mycolactone at the indicated times. Total virus was harvested at 24 hours and subjected to TCID50 titration on Vero cell. Data is presented as mean±SEM, n=3 biological repeats. Statistical significance comparing the mycolactone treatment to the DMSO control was assessed using one-way ANOVA and Dunnett’s multiple comparisons test. Statistical significance: p <0.01; **.

Taken together, these data reveal that while HSV1 sensitivity to Sec61 translocon blockade correlates with its kinetics of virus production, all three RNA viruses tested – RSV, ZIKV and in particular IAV - lose their sensitivity to mycolactone ahead of the production of infectious virus. This suggests that the threshold for virus envelope protein production is achieved early in infection by these viruses, and once that threshold has been reached, the ability to block virus production by targeting translocation of envelope proteins is greatly impaired.

## Discussion

A range of globally important human and animal virus pathogens are enclosed in an envelope which is derived from the host cell and contains virus-encoded membrane proteins, many of which are essential for virus infection. As such, the biogenesis of virus transmembrane proteins is a promising and broad-spectrum process for host-directed antiviral intervention of enveloped viruses and has the potential to be a first line of defence against novel and emerging virus pathogens.

There are various stages along the membrane protein biogenesis pathway that could be targeted, and there are examples of blocking virus production in culture by inhibition of onward trafficking from the ER using the drug brefeldin A, which disrupts the Arf1–COP1 coatamer system in the early secretory pathway [30-32], or targeting secretory pathway glycosylation enzymes to inhibit the maturation of envelope proteins [33-35]. Here we have chosen to target the process of Sec61-dependent protein translocation across the ER membrane using mycolactone, a potent inhibitor of this complex. We have studied this in a range of enveloped human viruses from different virus families and in different cell types to provide a comprehensive assessment of the broad-spectrum effects of mycolactone on virus infection.

The rationale for using mycolactone was based on its well-defined specificity for the Sec61 translocon and its relative potency, functioning as it does in the low nM range [16]. Our results have confirmed that the inhibitory effect of mycolactone on type I membrane proteins extends to a wide range of virus type I transmembrane proteins, whether they are translated from a standard transfection expression system or from viral mRNAs transcribed from their own genomes. Interestingly, expression of HSV1 gB was blocked in ectopically expressing cells but persisted at low levels in its non-glycosylated form in mycolactone-treated infected cells, indicating that the non-translocated gB may be stabilised by interaction with a viral binding partner during infection. The IAV type II transmembrane protein NA was susceptible to mycolactone during infection, in line with other studies on type II proteins [20], and although the RSV type II G protein persisted in its non-glycosylated form when produced by both ectopic and virus expression, *in vitro* translocation confirmed that mycolactone blocked its translocation across the ER membrane, while subcellular localisation assays indicated that it had not entered the secretory pathway for transport to the cell surface. Therefore, we propose a working model where translocation of G is blocked, but the mislocalised protein expressed in the cytosol is resistant to ubiquitination and/or proteasomal degradation. The SARS-CoV2 E protein, shown before to have type III topology [36],was resistant to mycolactone activity, as expected from studies on other type III proteins [19, 20]. Likewise, the two multipass proteins that were studied – gM from HSV1 and M from SARS-CoV2 – were both resistant to Sec61 blockade regardless of how they were expressed. Moreover, none of the gM protein that was produced in the presence of mycolactone was fully glycosylated, suggesting that the enzymes responsible for its modification are themselves blocked from entering the secretory pathway by mycolactone blockade of the Sec61 translocon, as shown elsewhere [22].

Despite all the RNA viruses that we tested producing transmembrane proteins that are essential for virus infectivity (S for SARS-CoV2; F for RSV; HA for IAV; E for ZIKV) there was a surprisingly short window during which mycolactone was able to fully block the production of progeny virus for the three of these viruses that we examined in more detail. Virus production of RSV, IAV and ZIKV continued even when mycolactone was added at a time before the initial rise in virus production, suggesting that sufficient envelope proteins were already in the system available for capsid envelopment to take place. For IAV, we were not able to find a time early enough after infection of MDCK cells where addition of mycolactone was able to block virus production, suggesting that synthesis of HA and NA occurs rapidly after virus entry. In the case of ZIKV, the translation of its full-length 380 kDa polyprotein is known to occur on the ER and involves the translocation of 18 TMDs across the ER membrane. Sec61 has previously been identified as a dependency factor for the replication of a number of other flaviviruses in a genome wide RNAi screen of insect and human cells [37], while inhibiting the Sec61 translocon in ZIKV infected cells with mycolactone blocked the formation of cytoplasmic replication vacuoles [38]. Our results presented here, where we have looked at expression of three cytoplasmic regions of the polyprotein (C, NS3 and NS5) confirm that this correlates with a lack of ZIKV polyprotein synthesis in mycolactone treated cells. Nonetheless, although early addition of mycolactone to ZIKV infected cells blocked virus replication, addition at a time just before virus production begins had only a limited effect on virus production, suggesting that as for the other RNA viruses, the full complement of structural proteins was available for morphogenesis ahead of the initiation of virus assembly.

HSV1 differed from the RNA viruses studied here in that mycolactone resulted in the immediate inhibition of virus production at the point of addition, suggesting that only the constituents that were available at that time could be utlilised for subsequent morphogenesis. HSV1 has a complex morphogenesis pathway which involves newly assembled capsids exiting the nucleus in a step termed nuclear egress. At around 100nm in diameter, HSV1 capsids are too large to move through nuclear pores and consequently they undergo a process known as envelopment-deenvelopment allowing the capsid to move through the inner and outer nuclear membrane [39]. Of note, a number of virus encoded membrane proteins are involved in nuclear egress - the UL34 type II membrane protein is known to be required for capsid to bud into the inner nuclear membrane [40] while the type I proteins gB and gH are proposed to be involved in fusion of the primary virion with the outer nuclear membrane resulting in the release of naked capsids into the cytoplasm where they are enveloped by endocytic membranes [41]. It is therefore likely that mycolactone not only inhibits transport of virus envelope proteins to the site of virus envelopment, but also inhibits the process of capsid egress from the nucleus, thereby blocking all steps in virus assembly. By comparison, although IAV also assembles its nucleocapsids in the nucleus, these are small enough to be actively exported through nuclear pores, a process which will proceed even in the presence of mycolactone allowing the capsids to continue with envelopment at the plasma membrane.

Intriguingly, virus production was rescued from the action of mycolactone in all infections at a time when virus envelope proteins were not yet detectable by Western blotting, suggesting that only small amounts of envelope proteins are required to produce significant amounts of progeny virions. This is in agreement with another of our studies on HSV1 where we found that a virus mutant which exhibits extreme translational shutoff of late proteins including virus envelope proteins nonetheless produced a similar level of progeny virus to wild-type HSV1 [28]. Our results presented here now suggest that enveloped viruses in general produce envelope proteins in vast excess to that required for virus maturation, raising the question of whether these high levels serve an additional purpose such as maintaining a favourable environment for virus survival, or counteracting some aspect of innate immunity.

Taken together, these studies demonstrate that virus systems offer a unique opportunity to assess mycolactone activity on membrane proteins as they are first produced. Moreover, the use of mycolactone has not only put a spotlight on the unexpected rapidity of virus morphogenesis but has also revealed its potential as a tool to study the cell biology of virus morphogenesis, to provide information about the timing, trafficking and targeting of virus proteins and the requirement for specific protein-protein interactions in virus assembly.

## Materials and Methods

### Cells and Viruses

HFFF, MDCK, Vero, HeLa, HaCaT, BSC1, Hek293T and HEp2 cells were cultured in Dulbecco’s modified Eagle medium (DMEM, Gibco) supplemented with 10% foetal bovine serum (FBS, Gibco) and 100 U/ml penicillin-streptomycin (Gibco). Hek293T cells expressing the mutant Sec61D60G protein [18] were routinely maintained in the presence of 200µg/ml hygromycin. The HSV-1 strain 17 (s17) used in this study was routinely propagated and titrated on Vero cells. The RSV A2 strain was kindly provided by Dalan Bailey (The Pirbright Institute) and was routinely propagated and titrated on HEp2 cells. The IAV Pr8 stain was kindly provided by Holly Shelton (The Pirbright Institute) and was routinely propagated and titrated on MDCK cells. The ZIKV MR766 strain was kindly provided by Kevin Maringer (The Pirbright Institute) and was routinely propagated and titrated on Vero cells. All viruses were propagated and titrated in DMEM supplemented with 2% FBS and 100 U/ml penicillin-streptomycin, with the exception of the IAV Pr8 strain which was propagated and titrated in DMEM supplemented with 1µg/ml TCPK-treated trypsin (Thermo Fisher Scientific) and 100 U/ml penicillin-streptomycin. Additionally, all MDCK cell monolayers were washed twice with PBS prior to infection with the IAV Pr8 strain.

### Antibodies and Reagents

Antibodies used in this study were kindly provided by the following individuals: gD, gB and gM, Helena Browne (University of Cambridge); gE, David Johnson (Oregon Health and Science University, Portland, OR USA); ZIKV NS3, and NS5, Kevin Maringer (The Pirbright Institute); SARS-CoV2 S, M, E and N, MRC-PPU Reagents, University of Dundee, UK. Other antibodies were purchased commercially: α-tubulin (Sigma), GFP (Clontech), Strep tag and giantin (AbCam); F for Western blotting (Santa Cruz Biosciences), F for immunofluorescence (Abcam), G (Invitrogen), (Abcam), IAV H1N1 (Bio-Rad), NP (Abcam), and ZIKV capsid (GeneTex). Synthetic mycolactone A/B was produced by Professor Yoshito Kishi (Harvard University, USA) [42]. Master stocks were prepared by dissolving the mycolactone to a concentration of 0.5 mg/ml in dimethylsulphoxide (DMSO, Sigma-Aldrich, St. Louis, MO, USA). Unless otherwise stated, cells were treated with a 31.25 ng/ml concentration of mycolactone and DMSO was used as a solvent control at 0.05%. Ipomoeassin F was a gift from Wei Shi (Ball State University) and was used at 250nM in transient transfection assays. Coibamide A was a kind gift from Kerry McPhail (Orgeon State University) and was used at 500nM in transient transfections and 1µM in in vitro translocation assays. PGNase F was purchased from New England Biolabs and was used as per manufacturer’s recommendations.

### Plasmids

For construction of pCDNA-gB and pCDNA-HA, the HSV-1 gB open reading frame (ORF) from wild type sc16 HSV-1 infectious DNA extracted from infected HaCaT cells was amplified with the primers TTGAAGCTTATGCGCCAGGGC (forward) and TTGGGATCCTTATTTACAACAAACCCCCCATC (reverse) and HA was amplified from the pHW2000_HA_PR8 plasmid (kindly provided by Holly Shelton, The Pirbright Institute) with the primers TGGAAGCTTATGAAGGCAAACCTACTGGT (forward) and TGGGGATCCTCAGATGCATATTCTGCACTGC (reverse) and inserted into pcDNA1.1/Amp. Additional plasmids used in this study were kindly provided by the following individuals: pcDNAgD and pcDNAgE, Helena Brown (University of Cambridge); pcDNA3-HSV1 gM, Colin Crump (University of Cambridge); pcDNAF, Monika Bajorek (French National Institute of Agricultural Research, France); Strep II tagged SARS-CoV2 N protein, M protein, E protein and S protein, Nevan Kroggan, (University of California, San Francisco, CA, USA); *in vitro* transcription/translation plasmids for SARS-CoV2 membrane proteins pcDNA5-S-OPG2, pcDNA5-OPG2-E and pcDNA5-M-OPG2, Stephen High (University of Manchester, UK). pEGFP-N1 plasmid was purchased from Clontech, pCMV-RSV-G(A-A2) was purchased from SinoBiological.

### Viral Infections

Cells seeded in 24-well plates were infected at a multiplicity between 0.2 and 2. The plates were incubated at 37°C for one (HSV-1, IAV, ZIKV, VACV) or two (RSV) hours to allow for virus adsorption, unless otherwise indicated. After adsorption, the inoculum was removed and cell monolayers were washed twice with PBS. After washing, the infection media was replaced and plates were returned to 37°C. If required, the infected cells were treated at the indicated times with either mycolactone or DMSO. Total virus was harvested for titration, or cells were lysed for Western blotting at the indicated times.

### Titration of HSV1 by Plaque Assay

Virus samples were serially diluted in the appropriate infection media. Six-well plates with confluent monolayers of cells were infected with 800µl of each of the different virus dilutions. The plates were incubated at 37°C for one hour to allow for virus adsorption. After adsorption, the inoculum was removed and cell monolayers were washed twice with PBS, and overlaid with fresh infection media containing 1% human serum (Sera Laboratories International Ltd). After three to four days, cells were fixed with formal saline and stained with crystal violet before plaques were counted. Viral titres were calculated and expressed as pfu/ml.

### TCID50 Assay

Cells were seeded into 96-well plates and virus samples were titrated onto cell monolayers. Briefly, virus samples were diluted 3-fold in infection media and cells were then inoculated with 200µl of each dilution in octuplicate. Inoculated cells were then incubated at 37°C for three (IAV), five (RSV) or seven (ZIKV) days, after which were fixed with 10% formal saline and stained with crystal violet before they were observed for the presence on cytopathic effect (CPE). Virus titres were calculated by the Reed and Muench method and were expressed as TCID50/ml.

### Cell Viability Assay

Cells were seeded into 96-well plates at a density that would give an 80-90-% confluent monolayer at the end of each assay. The cell monolayers were subjected to treatment with DMSO and varying concentrations of mycolactone diluted in DMEM supplemented with 10% FBS and 100 U/ml penicillin-streptomycin. For five-day assays, a 1:2 serial dilution of mycolactone was set up across the required rows of each plate resulting in final concentrations of 62.5 – 0.12ng/ml of mycolactone being used. For the 24-hour assays, cells were treated with two different concentrations of mycolactone as indicated. The plates were then incubated at 37°C for either five days or 24 hours. A resazurin solution (Sigma) was then added to wells (10% v/v) 4 to 8 hours (dependent upon the cell line) prior to the end of the assay and the cells were further incubated for the remaining duration at 37°C. The fluorescence of each plate at excitation 530nm and emission 558nm was then read using a CLARIOstar microplate reader. The cell viability percentage was calculated by normalising the absorbance readings from the cells treated with the different concentrations of mycolactone against the absorbance readings from the cells treated with DMSO.

### Transfections

HeLa cells seeded in 12-well plates were routinely transfected using 200 to 1000 ng plasmid DNA and 2 µl Lipofectamine™2000 (Invitrogen) according to the manufacturer’s recommendation and were simultaneously treated with either mycolactone or DMSO diluted in DMEM supplemented with 10% FBS and 100 U/ml penicillin-streptomycin. Twelve hours post-transfection, protein expression and localisation was assessed via Western blotting and immunofluorescence.

### SDS-PAGE and Western Blotting

Cells seeded in 12- or 24-well plates, were washed once with PBS and lysed in 1× SDS-lysis buffer at the indicated time points. Cell lysates were heated to 95 °C for 3 minutes and DNA sheared by pulling through a 23G needle. For gM western blots, samples were denatured at 42°C for 20 min to prevent aggregation of this transmembrane protein. For PGNase F treatment, cells were lysed in an NP-40 lysis buffer and were then treated with PGNase F according to the manufactures recommendation. Following treatment cell lysates were mixed in equal proportions with 2X SDS-lysis buffer. Proteins were separated on polyacrylamide gels ranging from 8% to 15% acrylamide and transferred onto nitrocellulose membranes. Membranes were blocked in 0.5% BSA or 5% milk in PBS-0.1% Tween (PBS-T) then incubated with primary antibody diluted in PBS-T. Following three washes in PBS 1% Triton X100, membranes were incubated with fluorescent conjugated secondary antibodies (Li-Cor Biosciences) diluted in blocking buffer. After three further washes in PBS-Triton, protein bands were visualized using an Odyssey CLx imaging system (Li-cor Biosciences).

### Immunofluorescence

Cells for immunofluorescence were grown on coverslips and fixed with 4% paraformaldehyde in PBS for 20 min. Fixed cells were permeabilized with 0.5% triton-X100 in PBS for 20 min and then blocked by incubation for 20 min in PBS containing 10% newborn calf serum. The cells were stained with primary antibody diluted in blocking buffer for 30 min and then washed extensively in PBS. The appropriate Alexa fluor conjugated secondary antibody (Invitrogen) diluted in block buffer was then added and the cells were incubated for a further 20 min. The coverslips were washed extensively in PBS and mounted in Vectashield containing DAPI (Vector Laboratories). Images were acquired using a 63× objective on a Nikon confocal microscope and processed using ImageJ 2.0 software and Photoshop.

### In vitro Translocation Assays

The pcDNAgD and pCMV-RSV-G(A-A2) plasmids encoding HSV-1 gD and the RSV attachment glycoprotein (G) respectively, were linearized then transcribed using the mMESSAGE mMACHINE™ T7 transcription Kit (Invitrogen). The mRNA was utilised within an Nuclease-treated Rabbit Reticulocyte System (Promega) *in vitro* translation assay. Briefly, the protein precursors were synthesised at 30 °C for 60 (HSV-1 gD) or 90 (RSV-G), supplemented with 8% v/v nuclease-treated canine microsomes (Dr Peter Mayerhofer, University of Surrey), 20 μM amino acids minus methionine (Promega), 20 U RNasin® Ribonuclease Inhibitor (Promega), EasyTag™ L-[35S]-Metionine (Perkin Elmer) and ∼1000 ng/μl of *in vitro* transcribed mRNA, in the absence or presence 200 ng/ml mycolactone or an equivalent volume of DMSO. Products produced using RSV G mRNA were layered onto a high-salt cushion and centrifuged at 100,000 g for 20 minutes using a Sorvall™ MTX 150 Micro-Ultracentrifuge and then the pellet was resuspended in low salt buffer.

All in vitro translation/ translocation assays conducted for SARS-CoV-2 proteins utilized pcDNA5 derived plasmids in a TnT® coupled transcription/translation system as per manufacturer’s instructions (Promega). Plasmid, 250ng/μl was utilised directly within the assay and all other conditions were followed as described above with the addition of 0.5μl of TNT RNA Polymerase T7. Reactions were incubated at 30°C for or 90 minutes and products were then layered onto a high-salt cushion and centrifuged at 100,000 g for 20 minutes and the pellet resuspended in low salt buffer. For endoglycosidase H (EndoH) treatment, an aliquot of the assay product was denatured in denaturing buffer (Promega) for 5 min at 95⁰C then cooled, diluted in Endo H Reaction Buffer (Promega) and incubated with 1000U Endo H (Promega) for 1hr at 37⁰C. All products were diluted with 2X SDS-PAGE lysis buffer and subjected to SDS-PAGE. The polyacrylamide gels were incubated for 15 minutes in Coomassie blue stain and then destained for 45 minutes. Prior to drying gels were incubated in a 10% v/v and 50% v/v ethanol solution to prevent cracking. The dried gels were exposed to a Phosphorimager screen and bands imaged on a Typhoon Phosphorimager (Typhoon) for 24-72hr.

### RT-qPCR

For the quantification of SARS-CoV-2 genome replication qRT-PCR (Thorne et al. 2020). Briefly, RNA was extracted from SARS-CoV-2 infected Caco-2 cells using the RNeasy Micro Kit (Qiagen) and residual genomic DNA was removed from the extracted RNA samples by on-column DNAse I treatment (Qiagen) as per manufacturer’s instructions. cDNA was then synthesized using SuperScript III with random hexamer primers (Invitrogen) and qRT-PCR was performed using a TaqMan Master mix (Thermo Fisher Scientific) using primers and a Taqman probe specific for SARS-CoV-2 E described elsewhere (Corman et al. 2020) on a QuantStudio 5 Real-Time PCR system (Thermo Fisher Scientific).

### Statistical Analysis

When required the data was normalised, and the DMSO and mycolactone treated cells were expressed as a percentage of the untreated cells (which is 100%). When multiple mycolactone conditions have been used, a one-way ANOVA was performed carried out with Dunnett’s correction for multiple comparisons relative to DMSO treated cells, which represents the solvent control. On the occasions where only one mycolactone treatment condition was used data was analysed by an unpaired two-tailed t test comparing the DSMO treated cells to the mycolactone treated cells. A four-parameter dose-response curve and variable slope model with the following equation: Y=Bottom + (Top-Bottom)/(1+10)^((LogIC50-X)*HillSlope)), was used to calculate IC50 values. All statistical analysis was performed using Graphpad Prism v8 software.

## Acknowledgements

The authors thank the following individuals for their generous provision of reagents used in this study: Kerry McPhail (Oregon State University, USA), Wei Shi (Ball State University, USA), Yoshito Kishi (Harvard University, USA), Dalan Bailey, Holly Shelton and Kevin Maringer (The Pirbright Institute, UK), Helena Browne and Colin Crump (University of Cambridge, UK), David Johnson (University of Oregan), Monika Bajorek (INRAE, Jouy-en-Josas, France), Nevan Kroggan (UCSF, USA), Stephen High (University of Manchester, UK), MRC-PPU Reagents (University of Dundee). We also thank Jane Newcombe and Peter Meyerhofer (University of Surrey) for technical assistance, and Sarah O’Keefe and Stephen High (University of Manchester) for advice on experiments.

LE and LK were funded by University of Surrey Faculty of Health and Medical Sciences PhD studentships. BH was funded by a Wellcome Trust Investigator Award to RS.

